# A missense mutation in zinc finger homeobox-3 (ZFHX3) impedes growth and alters metabolism and hypothalamic gene expression in mice

**DOI:** 10.1101/2022.05.25.493441

**Authors:** Patrick M. Nolan, Gareth Banks, Nora Bourbia, Ashleigh G. Wilcox, Liz Bentley, Lee Moir, Lee Kent, Rosie Hillier, Dana Wilson, Perry Barrett, Rebecca Dumbell

**Affiliations:** MRC Harwell Institute, Mammalian Genetics Unit and Mary Lyon Centre, Harwell Campus, Oxfordshire, UK; UK Health Security Agency, Centre for Radiation, Chemical and Environmental Hazards (UKHSA RCE), Harwell Campus, Oxfordshire, UK; The Rowett Institute, University of Aberdeen, Aberdeen, UK; Nottingham Trent University, School of Science and Technology, Clifton Lane, Nottingham, UK

**Keywords:** zfhx3, atbf1, energy balance, appetite, growth, mouse model, somatostatin, Gpr50, IGF1

## Abstract

A protein altering variant in the gene encoding zinc finger homeobox-3 (*ZFHX3*) has recently been associated with lower BMI in a human genome-wide association study. We investigated metabolic parameters in mice harbouring a missense mutation in *Zfhx3* (*Zfhx3*^*Sci/+*^) and looked for altered *in situ* expression of transcripts that are associated with energy balance in the hypothalamus to understand how ZFHX3 may influence growth and metabolic effects. One year old male and female *Zfhx3*^*Sci/+*^ mice weighed less, had shorter body length, reduced fat mass, smaller mesenteric fat depots, and lower circulating insulin, leptin, and insulin-like growth factor-1 (IGF1) concentrations than *Zfhx3*^*+/+*^ littermates. In a second cohort of 9 – 20-week-old males and females, *Zfhx3*^*Sci/+*^ mice ate less than wildtype controls, in proportion to body weight. In a third cohort of female-only *Zfhx3*^*Sci/+*^ and *Zfhx3*^*+/+*^ mice that underwent metabolic phenotyping from 6 - 14 weeks old, *Zfhx3*^*Sci/+*^ mice weighed less and had lower lean mass and energy expenditure, but fat mass didn’t differ. We detected increased expression of somatostatin, and decreased expression of growth hormone-releasing hormone and growth hormone-receptor mRNAs in the arcuate nucleus (ARC). Similarly, ARC expression of orexigenic neuropeptide Y was decreased and ventricular ependymal expression of orphan G protein-coupled receptor *Gpr50* was decreased. We demonstrate for the first time an energy balance effect of the *Zfhx3*^*Sci*^ mutation, likely by altering expression of key ARC neuropeptides to alter growth, food intake and energy expenditure.

## 1. Introduction

Obesity is a major contributor to preventable death worldwide (1), with comorbidities including type 2 diabetes, cardiovascular disease and some cancers, while early data even suggest a link to higher morbidity in COVID-19 patients (2). A genetic underpinning of body weight in humans has been demonstrated in twin and family studies (3), and genome wide association studies (GWAS) have identified over 250 common variants associated with obesity indicated by body mass index (BMI) in humans (4, 5). However very few of the mechanisms behind these associations have been described. Despite this, pathway analyses indicate that the majority of BMI – associated variants are in genes expressed in the hypothalamus (4), the brain region classically associated with homeostatic regulation of appetite and energy expenditure (6), indicating the importance of hypothalamic homeostatic body weight regulation in the human population. A recent GWAS identified a low frequency (4.34 %) protein altering variant in *ZFHX3* (zinc-finger homeobox-3) associated with lower BMI (−0.11 kg / m^2^), linking this gene to human obesity for the first time. In the same study, knockdown of the *Drosophila melanogaster* homologue (*zfh2*) in adipose and neuronal tissue led to increased triglyceride levels in these flies (7), implicating a conserved role for ZFHX3 and its homologues in body weight regulation.

ZFHX3, also called AT motif-binding factor-1 (ATBF1), was first described as a transcriptional regulator binding to an AT (adenine and thymine)-rich motif in the promotor of the *AFP* gene (8), and suppressing transcription by inhibiting binding of activator proteins (9). The mouse missense mutation “short circuit” (*Sci*) was identified in an *N*-ethyl-*N*-nitrosourea mutagenesis screen for circadian phenotypes and encodes a V1963F missense substitution in a highly conserved region in exon 9 of *Zfhx3* (10). *Zfhx3* is expressed in discrete regions of the adult mouse brain including nuclei of the anterior hypothalamus such as the suprachiasmatic nucleus (SCN) and arcuate nucleus (ARC) (11). Characterisation of the mouse missense mutant has implicated *Zfhx3* in SCN regulation of circadian rhythms. *Zfhx3*^*Sci/+*^ mice housed in constant darkness exhibit a shortened circadian behavioural rhythm believed to be mediated by altered ZFHX3 binding to AT motifs in the promoters of *Avp* and *Vip* in the SCN, although numerous other neuropeptide neurotransmitter and receptor levels were also misregulated (10). While important for the development of key brain regions (12), the circadian function of *Zfhx3* has been further demonstrated in a knockout of *Zfhx3* induced in adult mice (13), demonstrating a sustained role of this transcription factor in regulating neuroendocrine homeostasis, independent of any effects that may have been established throughout development. Despite the well-studied circadian effects of this mutation, growth and metabolic characterisation of *Zfhx3*^*Sci/+*^ mice has not previously been carried out.

RNA-seq analysis of hypothalamic tissue from *Zfhx3*^*Sci/+*^ mice has revealed altered expression of several genes implicated in appetite, growth and energy expenditure. Many of these identified transcripts, including those encoding somatostatin (*Sst*) and G-protein coupled receptor-50 (*Gpr50*), contain the AT motif in their promoter while ZFHX3-driven activity at such promoters is affected in the *Sci* mutation (10). This is a potential, although heretofore unexplored mechanism by which ZFHX3 may mediate whole body energy balance. Based on these previous findings, our study set out to investigate the role of ZFHX3 in hypothalamic transcriptional regulation of appetite, growth and energy expenditure pathways using aged and young adult *Zfhx3*^*Sci/+*^ mice. By comparing these pathways in control and mutant mice, we identify a potential mechanism for ZFHX3 regulation of growth, food intake, and energy expenditure through growth axis component mRNA expression in the ARC. This demonstrates mechanisms by which ZFHX3 may regulate appetite and body weight, identifying pathways that may be targeted for therapeutic intervention.

## 2. Methods

### 2.1 Animals

All animal experiments were conducted according to Medical Research Council (MRC) guidance in Responsibility in the Use of Animals for Medical Research (July 1993), and UK Home Office Project Licences 30/3206 and P6165EED1, with local ethical approval. All mice were housed at the Mary Lyon Centre, MRC Harwell Institute and, unless otherwise stated, were mixed-genotype group housed in individually ventilated cages in a pathogen free environment, in a 12h:12h light:dark light cycle (LD 12:12; lights on at 0700) with *ad libitum* access to standard rodent chow (SDS Rat and Mouse No. 3 Breeding diet, RM3), and water. *Zfhx3*^*Sci/+*^ mice, harbouring a V1963F missense mutation in the conserved region upstream of the 17th zinc-finger motif of ZFHX3, were maintained on a mixed C3H/HeH × C57BL6/J F1 background and genotyping was carried out by RT-PCR Taqman assay with the following primers and probes (5′ to 3′): FW TCCACGCATTGCTTCAGATG. REV TGTGCCTTCTGCTTGTTCTCA. Mutant probe VIC—TTTGAGCTCTTCATTCA. Wildtype probe FAM— CTTTGAGCTCGTCATT. Mice homozygous for this mutation show perinatal lethality (10) and so only mice heterozygous for this mutation and their wildtype littermates were included in this study.

#### 2.1.1 Experiment 1 – Year old mice

Male and female 12-month-old *Zfhx3*^*Sci/+*^ and *Zfhx3*^*+/+*^ mice (n = 9 – 12) underwent body composition analysis by quantitative NMR (Echo MRI, Echo Medical Systems, Houston, TX, USA), and the following week were fasted overnight, and humanely killed at 2 hours following lights on (zeitgeber time (ZT) 2) by overdose of anaesthesia. Under terminal anaesthesia, a retro-orbital bleed was carried out and serum collected, body length from nose to base of tail was measured and, on confirmation of death, left side fat tissues were dissected and weighed.

#### 2.1.2 Experiment 2 – Food intake measurements and early weight gain

Male and female *Zfhx3*^*Sci/+*^ and *Zfhx3*^*+/+*^ mice were weaned into pair-housed sex and genotype matched cages from 8 weeks of age. Mice and food remaining in the hopper were weighed weekly up to 20 weeks of age. For the food intake measurements, the experimental unit was the cage (n = 5 – 6 per sex and genotype), with n = 10 – 12 mice in total per group. Body weight was not collected for all mice at 19 – 20 weeks of age, so these data are excluded from analyses. Energy intake / body weight was calculated using combined body weight of mice pair-housed together at 15.21 MJ/Kg gross energy as specified for RM3 diet.

#### 2.1.3 Experiment 3 – Growth and hypothalamic gene expression in young female mice

Since early differences were stronger in female mice, only this sex was used in the final experiment. Female *Zfhx3*^*Sci/+*^ and *Zfhx3*^*+/+*^ mice were weighed and underwent body composition analysis by quantitative NMR (Echo MRI, Echo Medical Systems, Houston, TX, USA) every two weeks from 6 – 14 weeks of age (n = 24 – 26). 5 mice did not undergo measurement at week 14 so were excluded from the dataset, reducing sample size to n = 21 – 23. Overnight fasted glucose was measured at 8 weeks of age. At 10 weeks of age, urine was collected by scruffing the mouse over a microtube, if the mouse did not spontaneously urinate, this was encouraged by gently stroking the abdomen. At 12 weeks of age, indirect calorimetry was carried out using in PhenoMaster metabolic cages (TSE systems, Bad Homburg, Germany) for 48 h on a subset of mice (n = 16 – 19, see section 2.3 for details). Since gene expression may differ by time of day, we decided to collect tissues from these mice at two different time points, one in the light and one in the dark phase. Mice were then transferred to light controlled cabinets, where they remained group housed, and half of the mice underwent an advancement of light onset by 7 h so that there were now four groups of mice (*Zfhx3*^*Sci/+*^ and *Zfhx3*^*+/+*^, lights on 0700 and lights on 2400; n = 11 - 13). The mice were allowed to acclimate to the new 12 h : 12 h light : dark schedule for 14 days before they were humanely killed by cervical dislocation at ZT 3 or ZT 15 and the brain was collected and frozen on dry ice for storage at -70 °C.

### 2.2 Enzyme – Linked Immunosorbent Assays (ELISAs)

Leptin (Invitrogen, cat no KMC2281), insulin-like growth factor-1 (IGF1; R & D systems, cat no MG100), and insulin (Mercodia, cat no 10-1247) were measured in serum samples from mice in *experiment 1*, according to kit instructions. For the leptin ELISA, samples were diluted 1/10 and 1/20. Corticosterone was measured in urine collected from mice in *experiment 3* at ZT 1 and ZT 11, at either end of the light phase on the same day for each mouse. Only samples from mice with urine collected at both timepoints were measured (n = 15 – 16), according to kit instructions (Assaypro cat no EC3001-1) at a 1/20 dilution. For serum samples where measured values fell below the expected physiological range, and whose values were detected as potential outliers by Grubb’s test (P < 0.05), these were excluded from the results (Leptin: one female *Zfhx3*^*Sci/+*^, one male *Zfhx3*^*+/+*^; IGF1: one male and one female *Zfhx3*^*Sci/+*^, one female *Zfhx3*^*+/+*^). For urine corticosterone samples, several ZT 11 samples exceeded the limit of the standard curve, and it was not possible to remeasure, so these values were artificially capped at the maximum for that assay at 104.98 ng / µl.

### 2.3 Indirect Calorimetry

Mice were individually housed in PhenoMaster cages for 3 days. The respiratory exchange ratio, energy expenditure and food and water consumption were recorded throughout this period. However, to allow mice to acclimate to the novel environment, analysis of metabolic parameters was performed upon only the final 24 hours of recording. Food and water were available *ad libitum* throughout testing.

#### 2.4.1 Riboprobe synthesis

Riboprobes were previously synthesised for *Dio2, Dio3* (14), *Gpr50* (15), *Sst, Ghrh* (16), *Npy* (17), *Pomc* (18), *Trhr1* (19). For riboprobes not previously synthesised, a gene fragment 300-500 bp was synthesised with the following 5’ and 3’ overhangs containing M13 forward and reverse amplification sites and T3 and T7 polymerase sites respectively to facilitate amplification: 5’-GATTCGGAAACAGCTATGACCATGATTACGCCAAGCGCGCAATTAACCCTCACTAAAGGGAACAAAAGCTGG GTACC-3’, 5’-GAGCTCCAATTCGCCCTATAGTGAGTCGTATTACGCGCGCTCACTGGCCGTCGTTTTACAATT-3’ (gblock, IDT). New riboprobes were produced for *Bmal1* (accession no. NM_007489.4, range: 1389 – 1882), *Dlk1* (accession no. NM_010052.5, range: 337 – 610), *Ghr* (accession no. NM_010284.3, range: 1066 – 1556), *Otp* (accession no. NM_011021.5, range: 135 – 691), *Sim1* (accession no. U40575.1, range: 247 – 546), *Tbx3* (accession no. NM_011535.3 range: 1648 – 1963), and *Trh* (accession no. NM_009426.3, range: 114 – 463). Gene fragments were amplified by polymerase chain reaction amplification with M13 forward and reverse primers, which amplified 5’ upstream of polymerase sites. 100 ng of PCR product was used in an *in vitro* transcription reaction with T7 polymerase (antisense) in the presence of ^35^S-uridine 5 triphosphate (Perkin Elmer) for radioactive *in situ* hybridisation.

#### 2.4.2 *In situ* Hybridisation

Brains were cryosectioned on the coronal axis at 14 µm, mounted on poly-L-lysine coated slides (ThermoFisher) and stored at -70 °C. Radiolabelled i*n situ* hybridisation was performed as previously described (20): slides were fixed in 4 % PFA – 0.1M phosphate buffer before acetylation in 0.25 % acetic anhydride – 0.1 M triethanolamine, pH 8 and then ^35^S-labelled riboprobes (approx. 10^6^ counts / minute / slide) were applied to slides in hybridization buffer containing 0.3 M NaCl, 10 mM Tris-HCl (pH 8), 1 mM EDTA, 0.05 % tRNA, 10 mM dithiothreitol, 0.02 % ficoll, 0.02% polyvinylpyrrolidone, 0.02% bovine serum albumin and 10% dextran sulphate and hybridized for 16 h at 58 °C. Slides were then washed in 4x saline sodium citrate (SSC; 1x SSC: 0.15 M NaCl, 15 mM sodium citrate), incubated with RNase A (20 µg / µl) at 37 °C and washed in decreasing concentrations of SSC to 0.1x at room temperature and in a final incubation in 0.1x SSC at 60 °C. Slides were then dehydrated and opposed to Biomax MR film (Kodak) for 1-21 days as appropriate.

### 2.6 Statistical Analysis

Statistical comparisons were carried out using GraphPad Prism v 9.0.0. Data are expressed throughout as mean ± SEM and individual values plotted where appropriate. Comparisons were by unpaired t-test or 2-way ANOVA with Šídák’s multiple comparisons test. This was also carried out for continuous data when both sexes and genotypes were compared, with multiple 2-way ANOVAs being carried out to allow for pairwise comparisons within timepoints. For continuous data that only considered one sex, a 2-way repeated measures ANOVA was carried out followed by Šídák’s multiple comparisons test. For indirect calorimetry, 24 h, light phase and dark phase data were compared by lean mass-adjusted ANCOVA as recommended in (21), using quantitative NMR lean mass from 1 – 2 days before entering the phenomaster cages. When data did not meet the assumptions of the test, it was transformed and the analysis carried out on transformed data (*Sim1, Npy, Dio2;* log transformed), or non-parametric tests (Mann-Whitney and / or Wilcoxon matched-pairs signed rank) were used where appropriate and indicated in the text.

## 3. Results

### 3.1 *Zfhx3*^*Sci/+*^ mice are smaller and leaner than wildtype littermates

Since mice naturally gain weight with age, we first examined whole animal metabolic phenotypes in 12-month-old male and female *Zfhx3*^*Sci/+*^ mice (*experiment 1*). The *Sci* mutation led to a lower body weight in both male and female mice (Figure 1 A, P < 0.0001), and nose-base of tail length was significantly reduced in female mice and in *Zfhx3*^*Sci/+*^ mice, with those harbouring the *Sci* mutation having 0.87 ± 0.05 cm (8.27 ± 0.47 %) shorter body length than wildtype (Figure 1 B, Genotype: P < 0.0001, Sex: P = 0.0499, within sex both P < 0.0001). This was accompanied by lower lean mass in *Zfhx3*^*Sci/+*^ mice and female mice overall (Figure 1 C, Genotype: P < 0.0001, Sex: P < 0.0001, within sex both P < 0.0001). The same effect was seen for fat mass (Figure 1 E, Genotype: P = 0.0008, Sex: P < 0.0001, within female P = 0.0187), but when corrected to body mass, lean mass % and fat mass % only differed by sex overall (Figures 1 D, F; sex both: P < 0.0001). Since we were interested in whether the *Sci* mutation models the metabolic associations seen in humans, we calculated body weight / body mass^2^ which is similar to the human BMI measurement, and found this was significantly lower in *Zfhx3*^*Sci/+*^ mice (Figure 1 G; genotype P = 0.0178). On comparisons of fat depot mass (Figure 1 F – L), only mesenteric adipose tissue differed by genotype, with an overall reduction in *Zfhx3*^*Sci/+*^ mice (P = 0.0051), and interscapular brown adipose tissue approached a significantly lower mass in *Zfhx3*^*Sci/+*^ mice overall (P = 0.0535). However, when corrected for body weight, no effect of the *Sci* mutation was detected in any of the adipose depots measured (supplemental figure 1), so according to these measurements the adipose depots were proportionate to body weight in these smaller mice.

**Figure 1:**
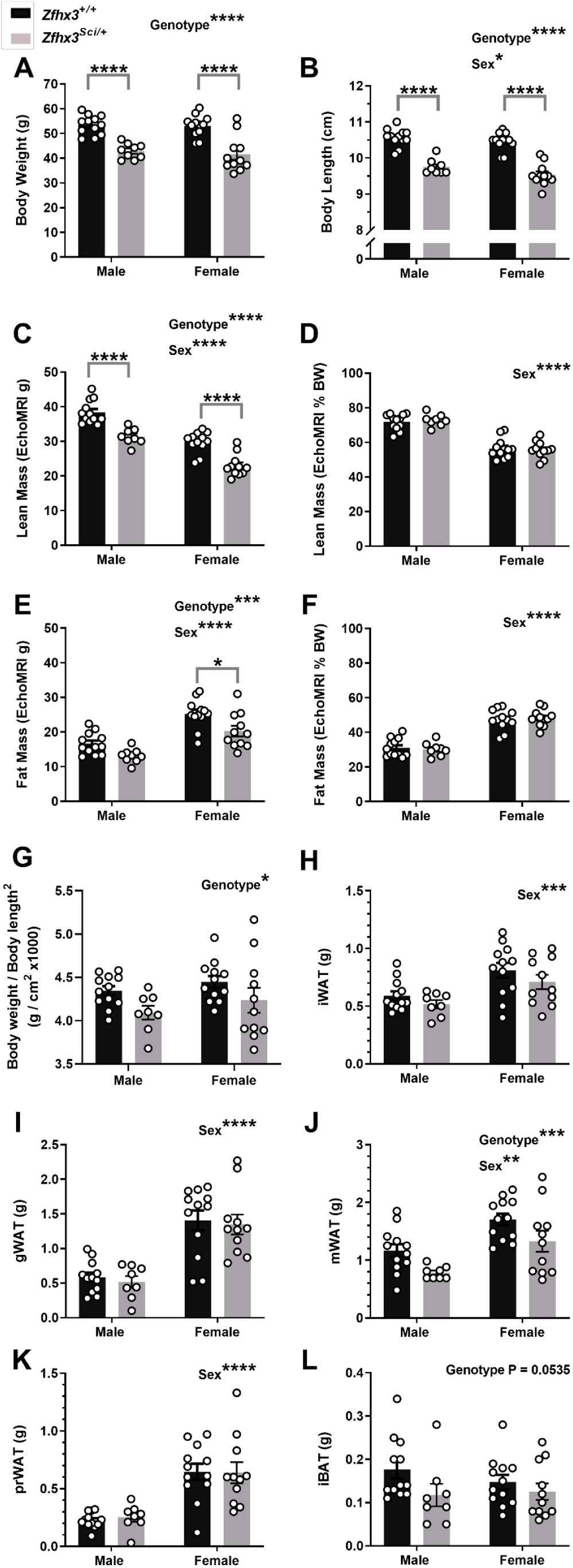
One year old *Zfhx3*^*Sci/+*^ mice are smaller than wildtype littermates. Male and female mice weigh less (A) and have shorter nose-tail body length (B) and lower lean (C) and fat mass (E) although this is proportionate to body weight (D, F). Body weight / body length^2^ was significantly lower in *Zfhx3*^*Sci/+*^ mice (G). Fat depot weights at dissection did not differ by genotype, except for mesenteric white adipose, which weighed less in *Zfhx3*^*Sci/+*^ mice (H-L). Plotted are mean ± SEM with individual values. Statistical comparison is by 2-way ANOVA, with overall results and within-sex comparisons indicated on the graphs. ****P<0.0001, ***P<0.001, **P<0.01, *P<0.05. BW: body weight, iWAT: inguinal white adipose tissue, gWAT: gonadal white adipose tissue, prWAT: perirenal white adipose tissue, mWAT: mesenteric white adipose tissue, iBAT: interscapular brown adipose tissue.

### 3.2 *Zfhx3*^*Sci/+*^ mice have lower circulating anabolic hormone concentrations

Comparison of fasted serum leptin revealed lower circulating levels overall in *Zfhx3*^*Sci/+*^ and female mice, with the genotype effect being stronger in the female values (Figure 2 A: Sex P < 0.0001, Genotype P = 0.0067, Sex x Genotype P = 0.0214, within female P = 0.0029), and when normalised to fat mass, only the sex difference remained (Figure 2 B; P < 0.0001), demonstrating that genotype effects on circulating leptin were likely a reflection of reduced adiposity in the *Zfhx3*^*Sci/+*^ mice. Fasted insulin was reduced in *Zfhx3*^*Sci/+*^ mice (Figure 2 C, P = 0.0014), suggesting a protective metabolic effect of the *Sci* mutation. Circulating IGF1 was also lower in *Zfhx3*^*Sci/+*^ mice (Figure 2 D, Genotype: P = 0.0004, Sex: P = 0.0152, within male P = 0.0118), implying, together with whole body mass and length differences, a somatic growth inhibition associated with the *Sci* mutation in 12-month-old mice.

**Figure 2:**
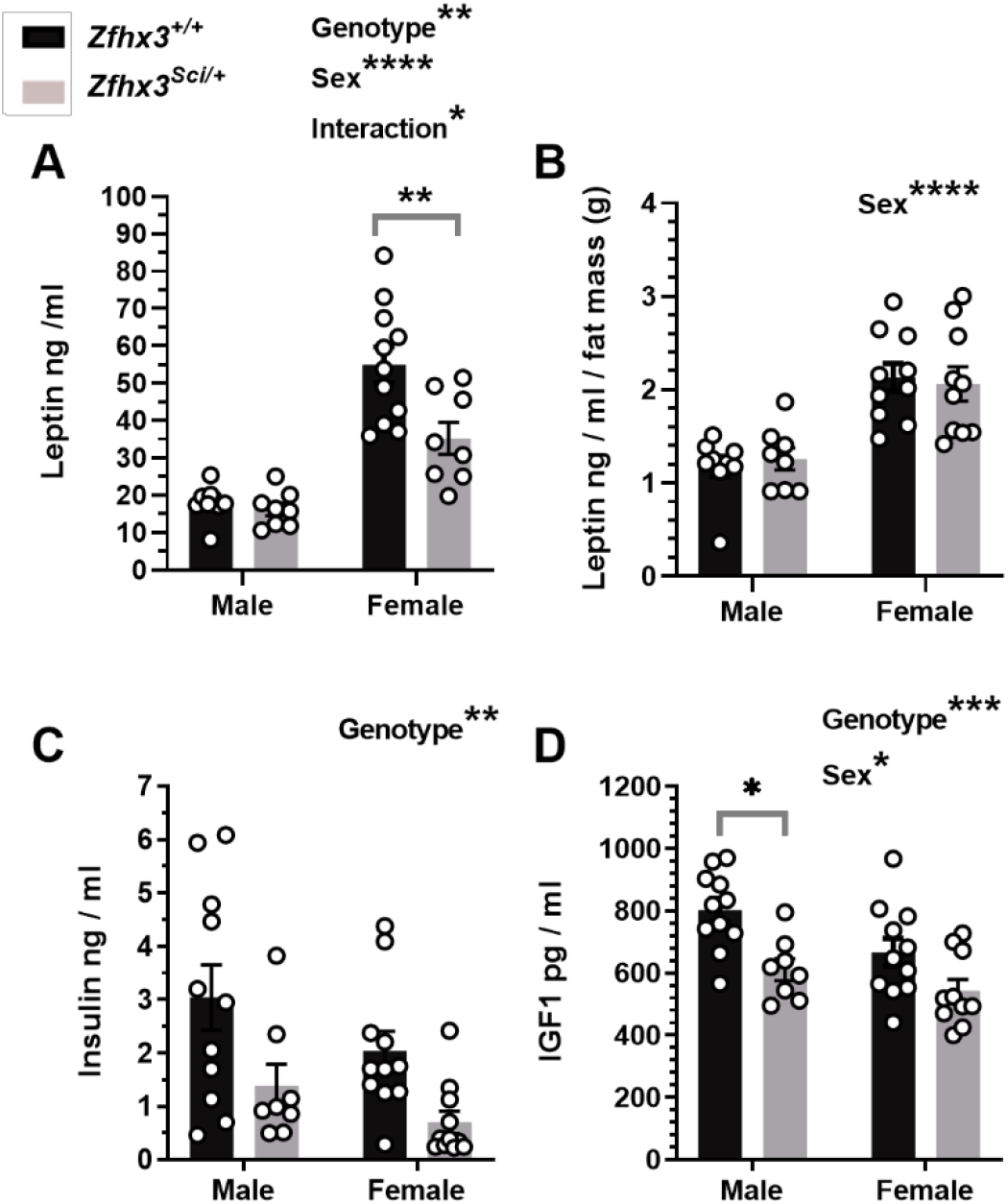
One year old *Zfhx3*^*Sci/+*^ mice have lower circulating concentrations of metabolic hormones. Circulating leptin was lower in fasted female *Zfhx3*^*Sci/+*^ mice (A), however this is proportionate to fat mass (g, EchoMRI; B). Circulating insulin and insulin-like growth factor-1 (IGF1) were also lower in *Zfhx3*^*Sci/+*^ mice. Plotted are mean ± SEM with individual values. Statistical comparison is by 2-way ANOVA, with overall results and within-sex comparisons indicated on the graphs. **** P < 0.0001, *** P < 0.001, ** P < 0.01, * P < 0.05.

### 3.3 *Zfhx3*^*Sci/+*^ mice consume less food than wildtype littermates, in proportion to body weight

To determine if the *Sci* mutation reduced food intake, a cohort of male and female *Zfhx3*^*Sci/+*^ and *Zfhx3*^*+/+*^ mice were pair-housed in genotype and sex matched cages and their food and body weight measured on a weekly basis from 8 – 20 weeks of age (*experiment 2*). Body weight was collected for all mice up to 18 weeks. In the second week of measurement, at 10 weeks of age, already a reduced cumulative food intake was detected in *Zfhx3*^*Sci/+*^ mice, and this was maintained through the continuation of the experiment (Figure 3 A, P < 0.01 weeks 10 -20), with an expected higher food intake in male mice compared to females detected in these same weeks (P < 0.001 weeks 10-20). Within-sex genotype differences were detected for several of these weeks for both male and female mice (weeks 10 – 14, 20 female, weeks 11 – 15, 20, male), with differences approaching significance for weeks 15 – 19 in female and 16 – 19 in male mice. Surprisingly, the *Zfhx3*^*Sci/+*^ body weight differences at this younger age were more apparent in female mice. As expected, female mice weighed significantly less than males throughout the experiment (Figure 3 B, week 8 P = 0.0355, week 9 P = 0.0007, weeks 10 – 18 P < 0.0001). *Zfhx3*^*Sci/+*^ mice weighed significantly less overall between weeks 9 – 13 (P < 0.05), and this difference approached significance for weeks 14 -18, however pairwise comparisons within female groups demonstrated a significantly lower body weight in *Zfhx3*^*Sci/+*^ mice from 10 weeks old onwards (P < 0.05), with interaction between sex and genotype reaching significance at weeks 15 and 18. When calculating the energy intake per g bodyweight for each cage of mice, no significant differences between sex or genotype were detected, indicating that mice ate in proportion to body weight.

**Figure 3:**
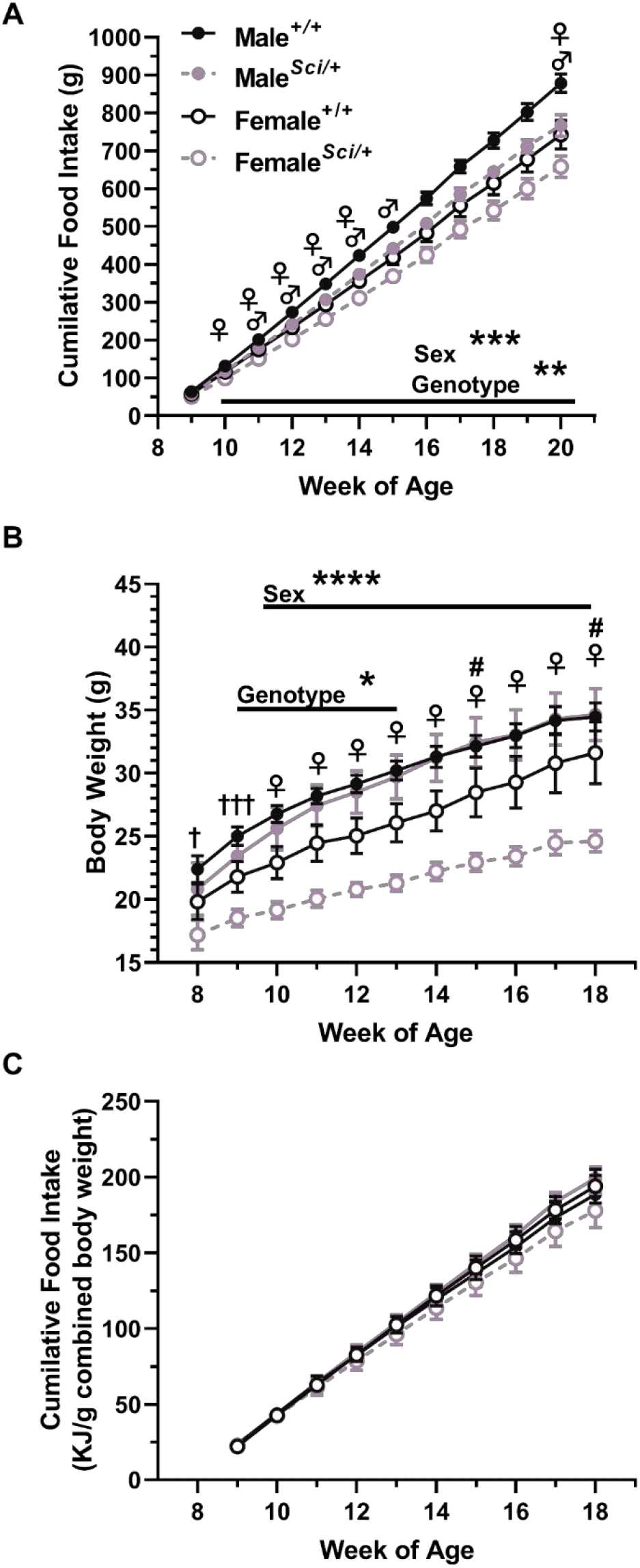
Male and female *Zfhx3*^*Sci/+*^ mice eat less than wildtype littermates. Mice were pair-housed (genotype and sex matched) and food weighed weekly (A; n = 5-6 cages). Male mice ate more than female mice from the second week of the experiment, and *Zfhx3*^*+/+*^ ate more than *Zfhx3*^*Sci/+*^ mice from the second week of the experiment. Weekly body weight revealed that the *Sci* mutation already inhibited weight gain from 9 weeks of age overall, with female *Zfhx3*^*Sci/+*^ mice weighing less than wildtype littermates from 10 weeks old onwards (B; n = 10-12). Although the genotype effect was not maintained from 14 weeks old onwards, a significant interaction could be detected at 15 and 18 weeks old. When energy intake was calculated per gram bodyweight (paired mouse weight) no significant differences were detected (C). Plotted are mean ± SEM, statistical comparison is by 2-way ANOVA per timepoint, and within-sex differences are indicated on the graph. **♀**P<0.05 *Zfhx3*^*Sci/+*^ vs *Zfhx3*^*+/+*^ female, ♂P<0.05 *Zfhx3*^*Sci/+*^ vs *Zfhx3*^*+/+*^ male, ***P<0.001, **P<0.01, † P<0.05 between sexes, † † † P<0.001 between sexes, # P<0.05 genotype – sex interaction.

### 3.4 Early differences in body weight are a result of impaired lean mass gain by *Zfhx3*^*Sci/+*^ female mice

To investigate more thoroughly early differences in weight gain, a cohort of *Zfhx3*^*Sci/+*^ and *Zfhx3*^*+/+*^ female mice were established, with the sample size established to humanely kill at two timepoints at the end of the experiment (*experiment 3*). Body weight, lean mass and fat mass were measured every two weeks from 6 – 14 weeks old. Body weight was lower in *Zfhx3*^*Sci/+*^ mice (Figure 4 A, P = 0.0002), with an interaction between time and genotype (P = 0.0470), reaching significance in pairwise comparisons from week 10. Lean mass was significantly lower in *Zfhx3*^*Sci/+*^ mice (Figure 4 B, P < 0.0001), and neither absolute fat mass nor % fat mass differed by genotype (Figures 4 C, D). Similarly, fasted glucose did not differ in *Zfhx3*^*Sci/+*^ mice at 8 weeks (supplementary Figure 2 A). As an indicator of peripheral and hormonal rhythms, urinary corticosterone was measured at timepoints approaching its expected peak and nadir levels (supplementary Figure 2 B). All mice had significantly higher urinary corticosterone at ZT 11 (Wilcoxon matched-pairs rank test, *Zfhx3*^*+/+*^ P = 0.0002, *Zfhx3*^*Sci/+*^ P < 0.0001), and no genotype differences were detected (Mann Whitney, NSD). To identify whether changes in metabolism underpinned the weight changes indicated above, a subset of mice underwent indirect calorimetry at 12 weeks of age. 24 h respiratory exchange ratio (RER) approached significantly lower levels in *Zfhx3*^*Sci/+*^ mice (Figure 4 E, G: genotype P = 0.0546; time x genotype P = 0.0048). RER was lower in *Zfhx3*^*Sci/+*^ mice when only the dark (active) phase was considered (P = 0.0306). There was a significant reduction in *Zfhx3*^*Sci/+*^ mice energy expenditure (time course, P = 0.0037), and an interaction between time and genotype (P < 0.0001), illustrated by significantly lower energy expenditure in *Zfhx3*^*Sci/+*^ mice in only the first 7 h of the dark (active) phase (Figure 4 F). This was confirmed when following correction for lean mass by ANCOVA, energy expenditure was significantly lower in *Zfhx3*^*Sci/+*^ mice during the dark phase (P = 0.0239), and total and light phase reductions approached significance (Figure 4 H). Food and water intake were measured during indirect calorimetry, and no genotype effect was detected in food intake over 24 h sample time (supplementary figure 2 C-D). However, a significant reduction in water intake was detected in *Zfhx3*^*Sci/+*^ mice (supplementary figure 2 E-F; Genotype P = 0.0125, Time x Genotype P = 0.0013; dark phase only P = 0.0020).

**Figure 4:**
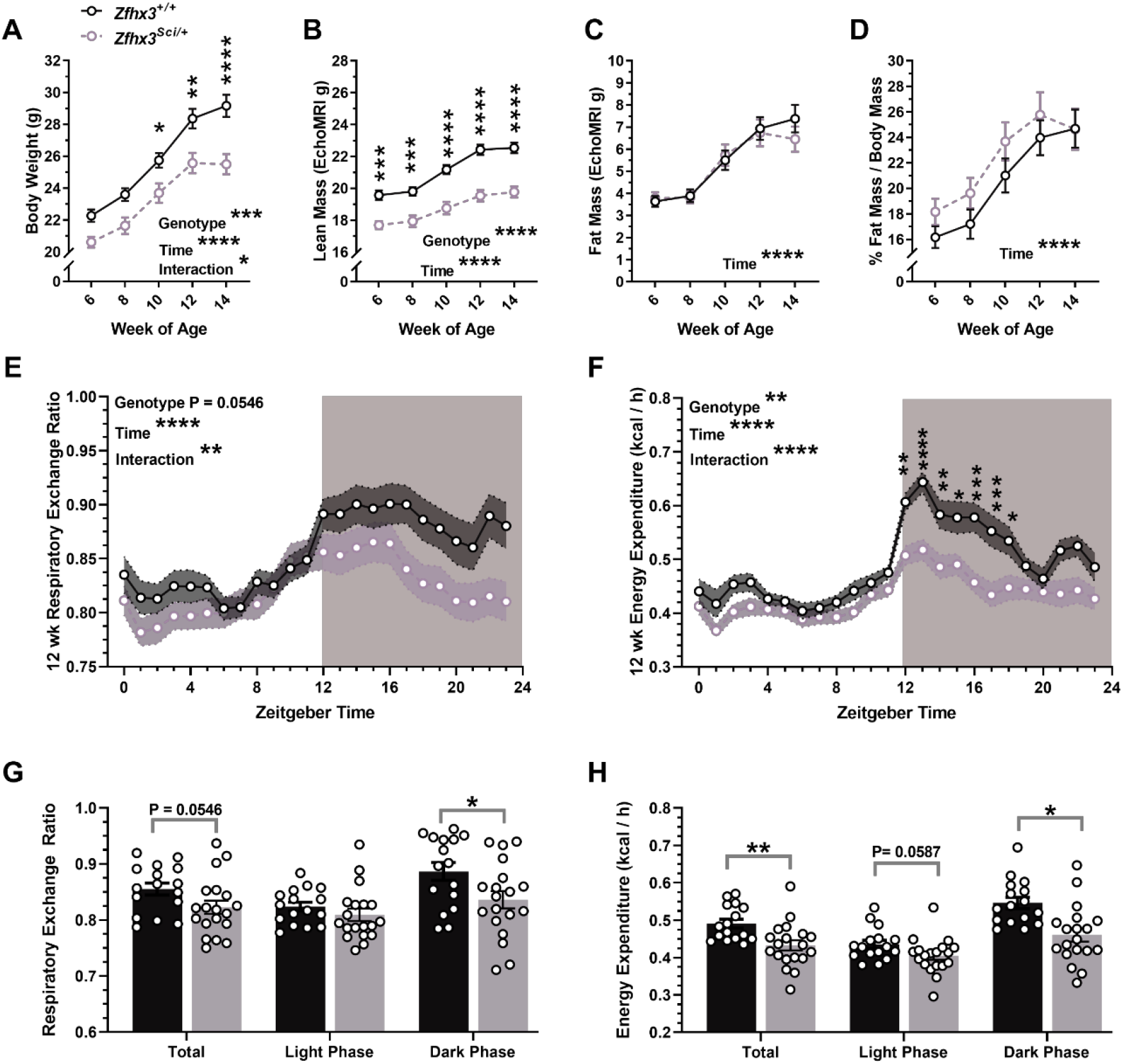
Early weight gain of female *Zfhx3*^*Sci/+*^ mice is lower than wildtype littermates, and driven by lower lean mass. The *Sci* mutation in female mice inhibits overall weight gain (A), composed of lean mass (B), but no differences in fat mass (C) or % fat mass (D). At 12 weeks old, these mice underwent indirect calorimetry and the second 24 h of measurements were compared to reveal a trend for lower 24 h RER and significantly reduced dark phase RER (E, G), while energy expenditure was reduced in *Zfhx3*^*Sci/+*^ mice (time course, F; lean mass ANCOVA adjusted, H). Plotted are mean ± SEM (A-H) with individual values (G-H). Statistical comparisons are by mixed effects model (REML) with Šídák’s multiple comparisons (A-D), RM-2-way ANOVA (E-F) and unpaired t-tests (G-H). A-D, n = 21 – 23; E-H n = 15 – 19. ****P<0.0001, ***P<0.001, **P<0.01, *P<0.05, NSD: no significant differences.

**Figure 4:** Early weight gain of female *Zfhx3*^*Sci/+*^ mice is lower than wildtype littermates and driven by lower lean mass. The *Sci* mutation in female mice inhibits overall weight gain (A), driven by lower lean mass (B), but no differences in fat mass (C) or % fat mass (D). Fasted blood glucose was not altered in these *Zfhx3*^*Sci/+*^ mice at 8 weeks old (E), and neither was urinary corticosterone at 10 weeks, and time of day differences were maintained (F). At 12 weeks old, these mice underwent indirect calorimetry and the second 24 h of measurements were compared to reveal a trend for lower 24 h RER and significantly reduced dark phase RER (G, I), while energy expenditure was reduced in *Zfhx3*^*Sci/+*^ mice (time course, H; lean mass ANCOVA adjusted, J). Plotted are mean ± SEM with individual values (E, F, I, J). Statistical comparisons are by mixed effects model (REML) with Šídák’s multiple comparisons (A-D), Mann-Whitney tests between genotypes (E, F), Wilcoxon matched – pairs tests (between timepoints, F), RM-2-way ANOVA (G, H), unpaired t-tests (I) and ANCOVA adjusting for lean mass (J). A-E, n = 21 – 23. **** P < 0.0001, *** P < 0.001, ** P < 0.01, * P < 0.05, NSD: no significant differences.

### 3.5 Hypothalamic mRNA expression is altered in female *Zfhx3*^*Sci/+*^ mice

Previous data suggests that *Zfhx3* may regulate hypothalamic gene expression. We hypothesised that such changes may be the cause of the metabolic effects we observed in *Zfhx3*^*Sci/+*^ mice. Therefore, candidate genes were investigated for altered expression in female *Zfhx3*^*Sci/+*^ mice in brains collected at ZT 3 in the light phase, and ZT 15 in the dark phase. These timepoints were chosen to reflect differential gene expression at different times of day. Genes were chosen based on previous observations as well as those associated with growth and energy balance in the hypothalamus. Table 1 summarises the genes investigated and where expression was detected. *Npy, Pomc, Sim1, Sst, Tbx3*, and *Trh* transcripts that had previously been implicated as altered by the *Sci* mutation in a previous RNA-seq experiment [10], were not detected in the SCN, indicating that expression of these transcripts may be altered in other hypothalamic nuclei.

**Table 1:** Brain regions investigated for mRNA expression. *In situ* hybridisation was carried out on coronal brain sections spanning the hypothalamus. Investigation for mRNA expression was carried out in different hypothalamic brain regions and quantified in figures 5 – 7. +: expression detected and quantified, -: expression not detected. SCN: suprachiasmatic nucleus, ARC: arcuate nucleus, PVN: paraventricular nucleus, PoPVN: posterior paraventricular nucleus, PeVN: periventricular nucleus, DMH: dorsomedial hypothalamus.

| Gene | Brain Region | mRNA detected? |
| --- | --- | --- |
| *Bmal1* | SCN | + |
| *Dio2* | Ventricular ependymal layer | + |
| *Dio3* | Ventricular ependymal layer | - |
| *Dlk1* | ARC | + |
| *Ghr* | SCN | - |
|  | PVN | + |
|  | ARC | + |
| *Ghrh* | SCN | - |
|  | PoPVN | + |
|  | ARC | + |
| *Gpr50* | SCN | - |
|  | Ventricular ependymal layer | + |
| *Npy* | SCN | - |
|  | ARC | + |
| *Otp* | SCN | - |
|  | PVN | + |
| *Pomc* | SCN | - |
|  | ARC | + |
| *Sim1* | SCN | - |
|  | PVN | + |
| *Sst* | SCN | - |
|  | PeVN | + |
|  | ARC | + |
| *Tbx3* | SCN | - |
|  | ARC | + |
| *Trh* | SCN | - |
|  | PVN | + |
|  | DMH | + |
| *Trhr1* | ARC | + |

#### 3.5.1. Hypothalamic growth axis component expression is altered in the arcuate nucleus

Radioactive *in situ* hybridisation was used to quantify spatially distinct expression of hypothalamic growth axis components. Expression of growth hormone-releasing hormone (*Ghrh*), the stimulatory arm of the growth axis, was decreased in the ARC of *Zfhx3*^*Sci/+*^ mice (Figure 5 A, P = 0.0088), with pairwise comparisons reaching significance in the dark phase (ZT 15, P = 0.0405). In the inhibitory arm of the growth axis, somatostatin (*Sst*) mRNA expression was increased in the ARC of *Zfhx3*^*Sci/+*^ mice (Figure 5 B, P = 0.0089), and ARC growth hormone-receptor (*Ghr*) expression was reduced overall in *Zfhx3*^*Sci/+*^ mice and with significantly increased overall expression at ZT 15 (Figure 5 C, Genotype P = 0.0333, ZT P = 0.0107). Interestingly, no differences in expression were detected in these genes in the more anterior hypothalamic nuclei (Figure 5 D: *Ghrh*, posterior hypothalamus (PoPVN), E: *Sst*, periventricular nucleus (PeVN), F: *Ghr*, paraventricular nucleus (PVN).

**Figure 5:**
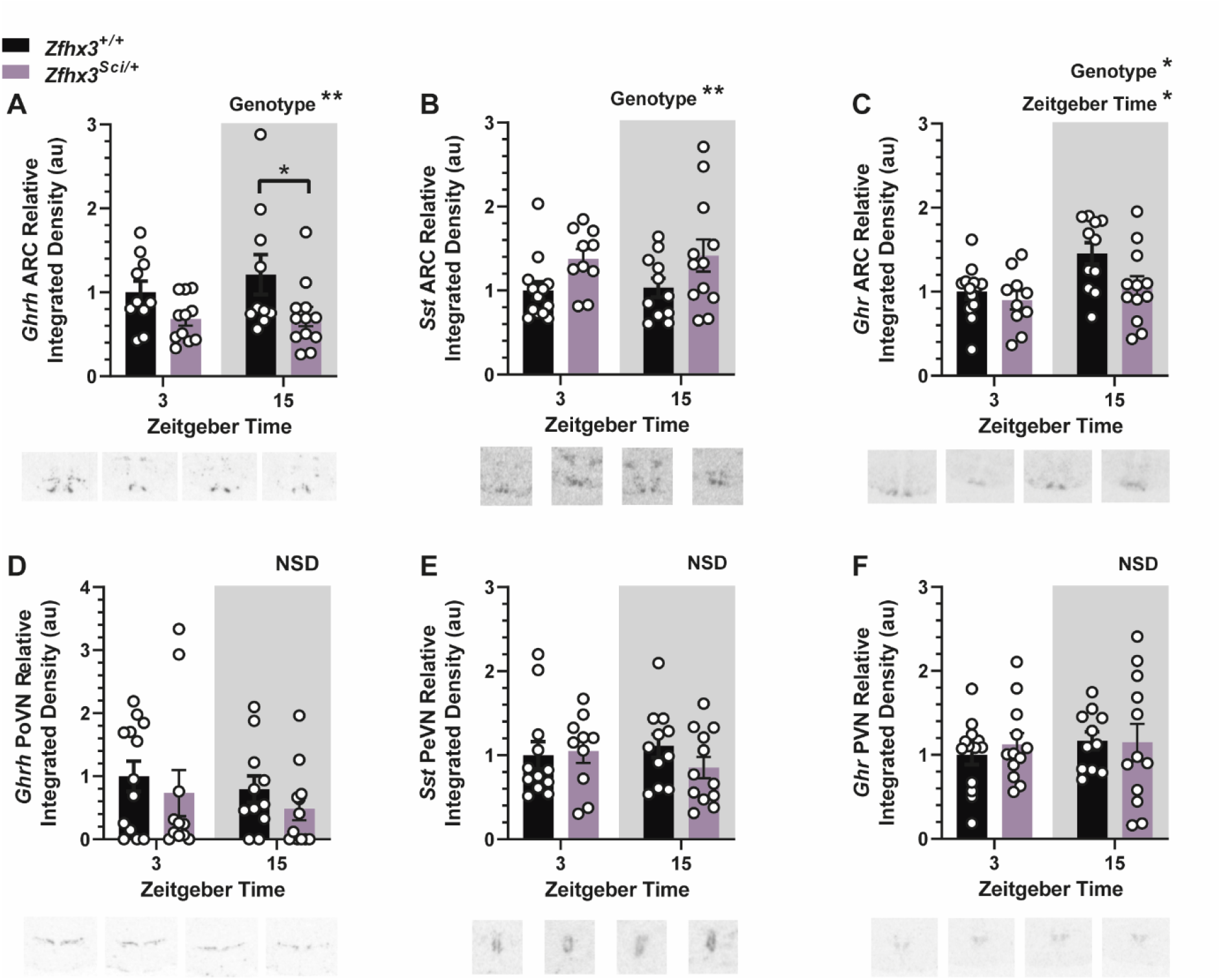
Hypothalamic expression of growth axis component genes is altered by the *Sci* mutation. Expression of growth hormone releasing hormone (*Ghrh*) is lower in *Zfhx3*^*Sci/+*^ mice arcuate nucleus (ARC, A) but not posterior paraventricular nucleus (PoPVN, D). Expression of somatostatin (*Sst*) is increased in *Zfhx3*^*Sci/+*^ mice in the ARC (B) but not periventricular nucleus (PeVN, E). Growth hormone receptor (*Ghr*) expression was lower in *Zfhx3*^*Sci/+*^ mice and at zeitgeber time (ZT) 3 in the ARC (C) but expression did not differ between groups in the paraventricular nucleus (PVN, F). Plotted are mean ± SEM with individual values overlaid. Example images are shown beneath each plot. Comparisons are by 2-way ANOVA with Šídák’s multiple comparisons tests indicated where appropriate. * P < 0.05, ** P < 0.01, NSD: no significant differences. N = 10 – 12.

#### 3.5.2 Additional candidate hypothalamic gene expression altered by the *Sci* mutation

We measured expression of the circadian gene *Bmal1* as an indicator of rhythmic gene expression in the suprachiasmatic nucleus (SCN). Overall, there was a significantly higher *Bmal1* expression in the dark phase (Figure 6 A, P = 0.0008), and an interaction between ZT and genotype (P = 0.0052). Multiple comparisons indicated increased *Bmal1* expression in the dark phase only for *Zfhx3*^*+/+*^ mice (P = 0.0002) and lower expression in *Zfhx3*^*Sci/+*^ mice at ZT 15 (P = 0.0445). This may indicate dysregulation of *Bmal1* circadian expression. Expression of appetite suppressing peptide *Pomc* approached a significant reduction in the ARC of *Zfhx3*^*Sci/+*^ mice (Figure 6 B, P = 0.0607) and appetite stimulating peptide *Npy* was significantly reduced overall in *Zfhx3*^*Sci/+*^ mice, with a significantly lower expression in *Zfhx3*^*Sci/+*^ mice at ZT 15 (Figure 6 C, Genotype P = 0.0294, within ZT 15 P = 0.0365). Expression of *Gpr50* in the ventricular ependymal layer was overall increased in *Zfhx3*^*Sci/+*^ mice, reaching significance in pairwise comparisons at ZT 15 (Figure 6 D, Genotype P = 0.0188, within ZT 15 P = 0.0325). Further genes investigated for altered expression did not differ with genotype (supplementary Figure 4). Expression of *Otp* in the paraventricular nucleus (PVN) was reduced overall at ZT 3 (supplementary Figure 4 A, P = 0.0015), and expression of ventricular ependymal layer *Dio2* was increased at ZT 15 (supplementary Figure 4 D, P = 0.0285) as was dorsomedial hypothalamus (DMH) *Trh* (supplementary Figure 4 F, P = 0.0145). Expression of PVN *Sim1, Trh*, ARC *Trh-r1, Dlk1* and *Tbx3* did not differ by genotype or ZT (supplementary Figure 4).

**Figure 6:**
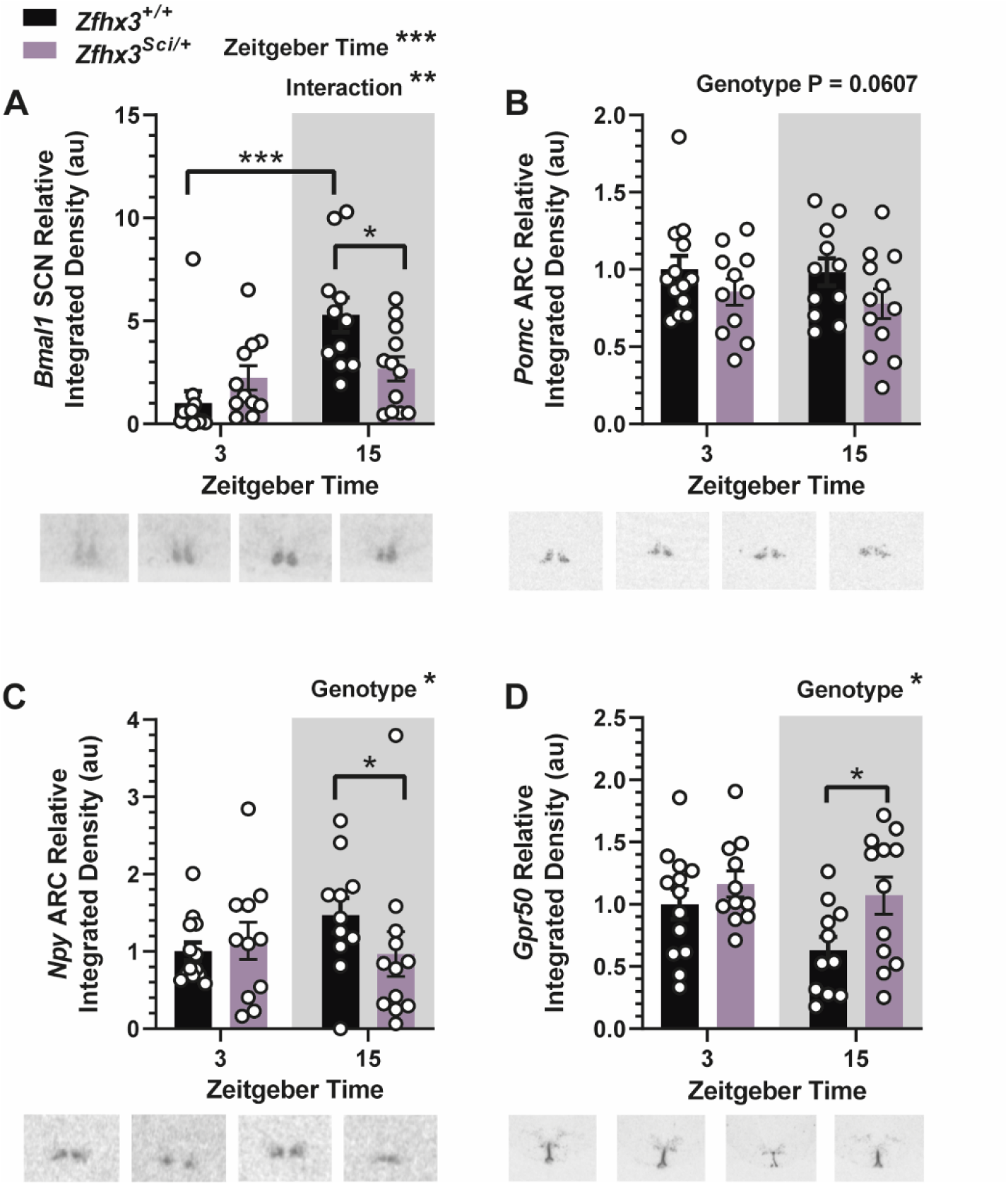
Additional hypothalamic gene expression is altered by the *Sci* mutation. Expression of *Bmal1* in the suprachiasmatic nucleus (SCN) was increased at ZT 15 overall, with an interaction between ZT and genotype, and reduced expression in *Zfhx3*^*Sci/+*^ mice at ZT 15 only (A). Expression of *Pomc* in the arcuate nucleus (ARC) approached a significant reduction in *Zfhx3*^*Sci/+*^ mice (B), and ARC expression of *Npy* was overall reduced in *Zfhx3*^*Sci/+*^ mice (C). Expression of *Gpr50* was increased overall in the ventricular ependymal layer, with pairwise comparisons reaching significance at ZT 15. Plotted are mean ± SEM with individual values overlaid. Representative images are shown beneath each plot. Comparisons are by 2-way ANOVA with Šídák’s multiple comparison tests indicated where appropriate. *Npy* data were log transformed for statistical comparison. * P < 0.05, ** P < 0.01, *** P < 0.001. NSD: no significant differences. N = 10 – 12.

## 4. Discussion

This study establishes an energy balance phenotype in *Zfxh3*^*Sci/+*^ mice aged to 12 months old, with younger mice demonstrating a slower increase in lean mass from the earliest measured timepoint at 6 weeks of age. These data are therefore consistent with a progressive effect of inhibited growth and food intake over the lifespan of *Zfxh3*^*Sci/+*^ mice. This was accompanied by altered hypothalamic gene expression in growth and appetite regulating peptide mRNAs in the ARC.

We demonstrate that both male and female *Zfxh3*^*Sci/+*^ mice weigh less and have lower adiposity at 1 year. Although reduced fat mass was accounted for by smaller body size, and lower circulating fasted leptin was in proportion to fat mass, these mice ate less and had a lower fasted insulin level. Notably, there was low circulating IGF1 in *Zfxh3*^*Sci/+*^ mice, an indicator of growth axis suppression. Indeed, the overall phenotype is reminiscent of a GH activity deficiency phenotype (22). IGF1 null mice have low viability and are born small, with inhibited growth (23). Liver specific deletion of IGF1 leads to elevated levels of growth hormone and this is presumed to contribute to the insulin resistance but not elevated blood glucose they experience (24). Since we were unfortunately unable to measure growth hormone in the present study, it is assumed that reduced IGF1 was a consequence of lowered growth axis tone, likely driven from the altered gene expression in the hypothalamus, and not a direct effect of the *Zfhx3*^*Sci*^ mutation, however this cannot be discounted. The reduced somatic growth effect was demonstrated in female mice from 6 weeks of age, when we already detected lower lean and whole-body mass. It is tempting to extrapolate from this data that a fat mass gain was starting to wane for *Zfxh3*^*Sci/+*^ mice at 16 weeks, leading to the fat mass differences seen in the aged mice, but these measurements were not possible given the constraints of the experimental plan.

It is reasonable to consider whether the phenotype observed in *Zfxh3*^*Sci/+*^ mice is truly a metabolically protective phenotype, or merely models short stature. Indeed, in a recent human exome sequencing study *ZFHX3* was identified as a candidate gene for short stature, interacting with growth hormone pathway component *POU1F1* (25). However multiple studies have found the growth axis to be intrinsically linked with energy balance (26), and when calculating the relationship between body length and weight in *Zfxh3*^*Sci/+*^ mice, a measure similar to BMI in humans, these mice still had a lower index than wildtypes. While BMI is arguably a poor measure of adiposity or metabolic dysfunction, the comparison we demonstrate indicates that the *Zfxh3*^*Sci/+*^ mice model the phenotype of people harbouring the human Q2021H *ZFHX3* variant (7).

*Sst* and *Gpr50* contain the AT motif that ZFHX3 can bind to alter expression (10), and therefore likely drive the observed phenotype. When considering changes in growth axis component gene expression in the hypothalamus of *Zfxh3*^*Sci/+*^ mice, it was perhaps unsurprising that expression was altered only in the ARC, where *Zfhx3* is demonstrably expressed (11), with the inhibitory arm (somatostatin) upregulated and the stimulatory arm (GHRH) downregulated. Somatostatin is released from neurons in the periventricular nucleus to inhibit growth hormone release from the pituitary and do not originate in the ARC. Therefore, ARC *Sst* neurons likely have a different role. It has long been shown that the *Sst* neurons in the ARC predominantly project within the ARC to *Ghrh* neurons (27, 28), which then project to the median eminence and release GHRH to somatotrophs of the anterior pituitary to alter growth hormone secretion (29). More recent work in human tissue supports active synapses from somatostatin to GHRH neurons within the ARC (30). Thus, although we cannot rule out direct regulation by ZFHX3 of GHRH, the parsimonious explanation is that altered ZFHX3 binding to the *Sst* AT motif in *Zfxh3*^*Sci/+*^ mice leads to upregulation of somatostatin, with subsequent inhibition of *Ghrh* production and suppression of the growth axis. We have previously demonstrated that somatostatin analogues inhibit somatic growth and circulating IGF1 in Siberian hamsters (*Phodopus sungorus*) (31, 32), driving changes in body weight and fasting glucose and insulin as well as increased incidence of daily torpor (33). This supports a conserved role for these ARC *Sst* neurons across mammalian species.

Lower food intake in *Zfxh3*^*Sci/+*^ mice was in proportion to body weight, and the experiments we conducted were not able to determine whether food intake influenced body mass or vice versa, although this would be interesting to explore in future work. We found suppressed food intake in both male and female *Zfxh3*^*Sci/+*^ mice from 10 weeks old and detected decreased expression of the orexigenic peptide *Npy* mRNA in 14-week-old female mice. Links between NPY and the growth axis are well established (26), stimulating GHRH and suppressing somatostatin release from PeVN neurons to the hypophyseal blood portal system to reach pituitary somatotrophs in sheep (34, 35), and genetically altered mice (36). In broiler chickens, treatment with growth hormone decreases NPY levels (37), and *npy* expression is reduced in zebrafish (*Danio rerio*) overexpressing growth hormone (38). Intracerebroventricular (ICV) injection of NPY decreases plasma growth hormone in rats, postulated to be via stimulation of somatostatin neurons (39, 40), and ICV injection of growth hormone increases *Npy* and *Agrp* mRNA in the ARC (41). Following hypophysectomy, low *Npy* expression in the ARC can be rescued, and stimulation of *c-fos* is seen in NPY neurons of the ARC with chronic treatment with growth hormone (42, 43), and mice lacking the growth hormone-receptor in AgRP/NPY neurons are protected from hypothalamic changes induced by food restriction (41). Notably, much less is known about interactions between NPY and somatostatin neurons of the ARC. Due to the lack of known ZHFX3-binding motifs in *Npy*, it is not clear whether the *Sci* mutation directly alters expression of this gene, whether altered expression is a downstream effect of the suppressed somatic growth axis, or via some other mechanism.

We found that energy expenditure was lower in *Zfxh3*^*Sci/+*^ mice, particularly in the dark phase when mice are usually more active. We also found a higher expression of *Gpr50* mRNA in the ventricular ependymal layer of the hypothalamus in the same mice. The mouse *Gpr50* promotor contains an AT motif that ZFHX3 binds to alter expression (10). This orphan G protein-coupled receptor is implicated in regulation of energy expenditure; mice lacking *Gpr50* weigh less and are protected from diet induced obesity, likely due to increased energy expenditure (44). It is therefore possible that increased expression of *Gpr50* contributed to the lower energy expenditure seen in *Zfxh3*^*Sci/+*^ mice.

The use of GWAS to identify genetic drivers of obesity and therefore potential targets for intervention can be controversial, as most common variants are in non-coding regions of the genome and thus present significant challenges to investigation and interpretation. The first time ZFHX3 was associated with human body weight was when a rare protein altering variant was identified in a human population of predominantly European ancestry (7). This Q2021H variant is in a similarly highly conserved region to the *Sci* mutation and knockout of the *Drosophila* homologue (*zfh2*) led to increased whole-body triglycerides, which the authors interpreted as increased adiposity, whether knocked down in neural or adipose tissue in the flies. Considering the phenotype described in the present study, it is tempting to hypothesise that the low frequency human Q2021H ZFHX3 variant, present in 4.34 % of sampled populations, acts through a similar pathway to that seen in the V1963F *Sci* mouse mutation.

## 5. Conclusion

For the first time, we demonstrate a role for ZFHX3 to alter growth and energy balance in a mouse with a protein altering mutation similar to a variant found at low frequency in the human population, both in a highly conserved region of the gene encoding this transcription factor. We have shown that this *Sci* mutation conveys lower food intake and altered hypothalamic gene expression, along with lower fat mass and individual depot weights in proportion to overall body mass, and lower fasted IGF1, leptin and insulin levels. This suggests that this small body size phenotype may convey protection from metabolic disease. While investigating altered hypothalamic gene expression using *in situ* mRNA measurements, we identified several transcripts with altered expression in the ARC and in the ependymal layer of the third ventricle of the hypothalamus that may drive this small body size phenotype. This implicates ARC growth axis components *Sst, Ghrh Ghr* and *Npy;* and ventricular ependymal layer *Gpr50* in driving small body size, lower food intake and altered energy expenditure in *Zfhx3*^*Sci/+*^ mice. This may together explain the association between the human Q2021H variant with lower BMI. Further work outside the scope of this investigation will be able to determine further the specific role of these identified pathways to influence energy balance and better explain the role of ZFHX3 and the central growth axis in body weight regulation.

## Appendices

Supplementary figures are provided in a separate file.

## Acknowledgements

The authors thank the staff of the Mary Lyon Centre and core services at MRC Harwell Institute for assistance with mouse studies. PMN was supported by the Medical Research Council (MC_U142684173), the Barrett lab was supported by the Scottish Government, R. Dumbell was supported by the Medical Research Council (Roger Cox: MC_U142661184) and Nottingham Trent University as well as a Research Visit Grant from the British Society for Neuroendocrinology to carry out this work. A version of this manuscript was previously uploaded to the bioRxiv preprint server (45)

## Author Contributions

RD, PMN, NB and PB contributed to conceptualisation of the project; mouse colony management was carried out by RH and LK; experiments and sample collection were performed by RD, GB, AGW, LB, LM, LK and DW; the original draft was written by RD and reviewed and edited by RD, PMN, GB and PB, with support from all authors.

