## Supplementary Figure for "A missense mutation in zinc finger homeobox-3 (ZFHX3) impedes growth and alters metabolism and hypothalamic gene expression in mice"

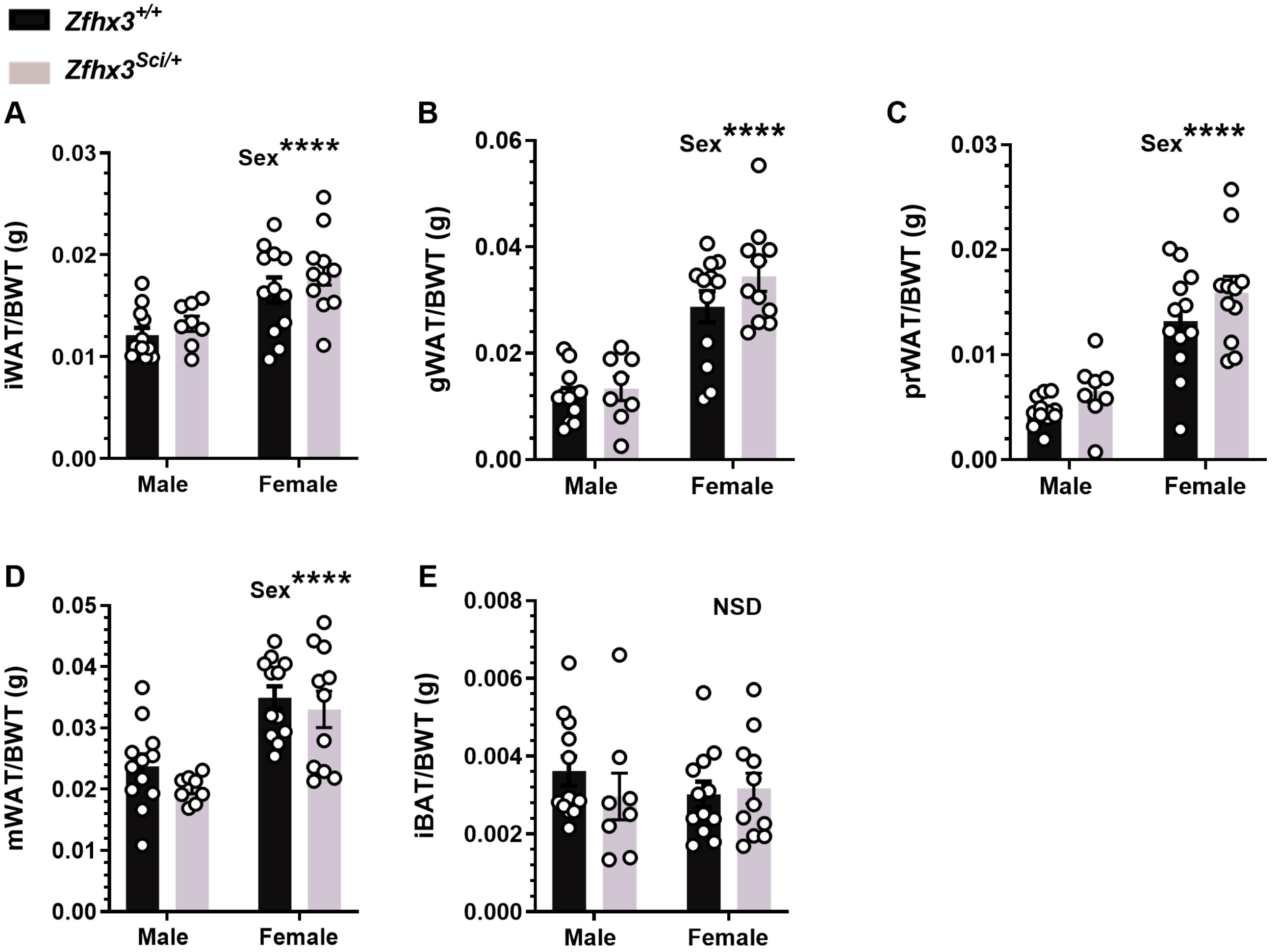


**Supplemental Figure 1: One year old *Zfhx3$^{Sci/+}$* mice fat pad mass do not differ to wildtype when corrected for body weight.**

The *Sci* mutation in male and female mice does not alter fat pad mass / body weight (BWT). This corrected tissue mass is higher overall in female mice in al white adipose tissues measured (A-D), and sex did not affect brown adipose corrected mass (E). Plotted are mean ± SEM with individual values. Statistical comparison is by 2-way ANOVA, with overall comparisons indicated on the graphs. ****P<0.0001, NSD: no significant differences. iWAT: inguinal white adipose tissue, gWAT: gonadal white adipose tissue, prWAT: perirenal white adipose tissue, mWAT: mesenteric white adipose tissue, iBAT: interscapular brown adipose tissue.

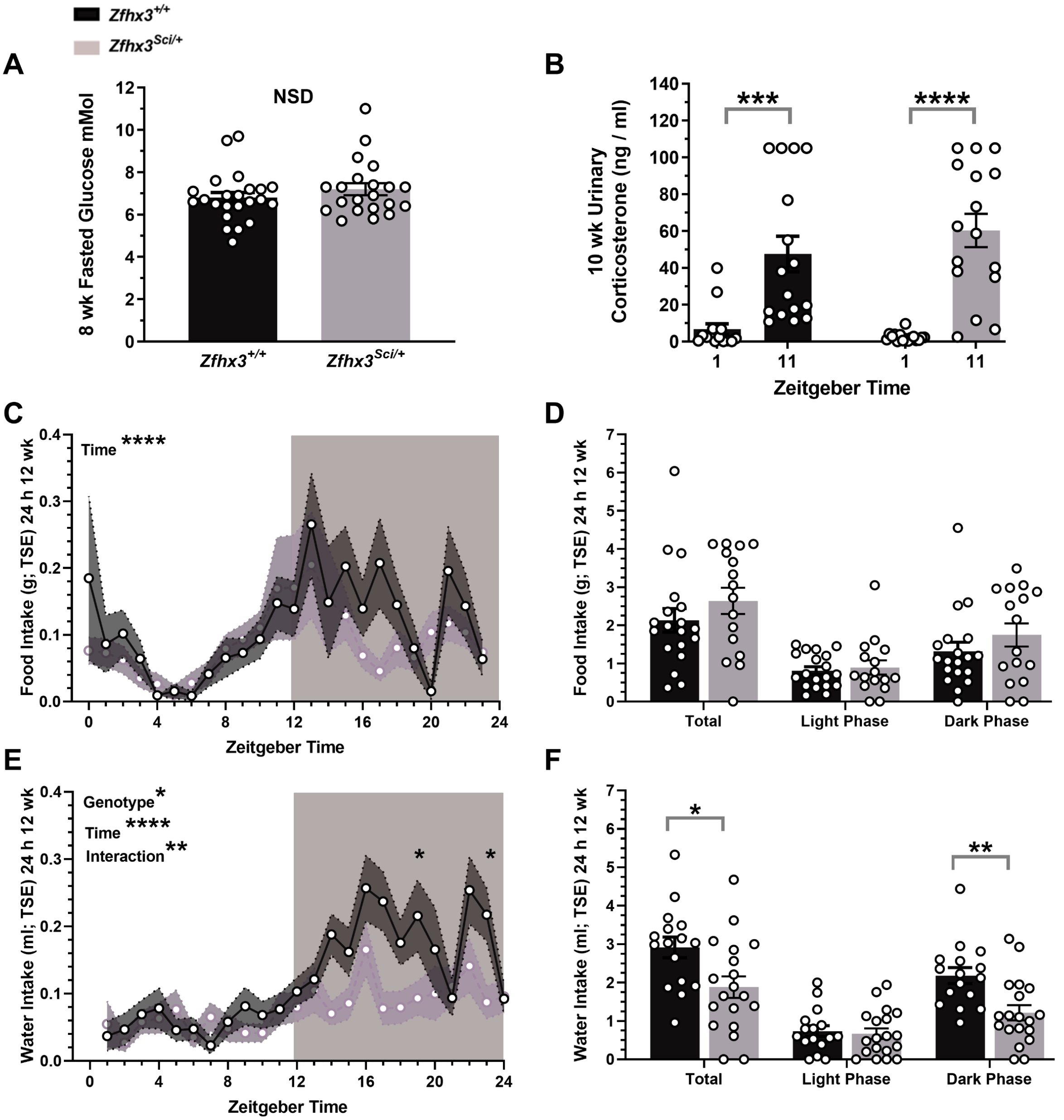


**Supplemental Figure 2: 8 week blood glucose, 10 week urinary corticosterone and 24 h food and water intake data from TSE Phenomaster Calorimetry in 12 week old female *Zfhx3$^{Sci/+}$* mice.**

Fasted blood glucose was not altered in *Zfhx3$^{Sci/+}$* mice at 8 weeks old (E), and neither was urinary corticosterone at 10 weeks, with peak and trough values maintained (F). In the TSE phenomaster data experiments, food intake in 1 h bins (C) total intake over 24 h, and light and dark phase only (D) was not altered by genotype during the sampling period in TSE Phenomaster metabolic cages, carried out in the second 24 h single housed in metabolic cages, at 12 weeks of age. However, water intake was significantly reduced overall in *Zfhx3$^{Sci/+}$* mice, with an interaction between time and genotype (E) since water intake was significantly reduced in the dark phase and not the light phase (F). Plotted are mean ± SEM with (A, B, D, F) or without (C, E) individual values. Statistical comparison is by Mann-Whitney (A,B), Wilcoxon matched – pairs tests (between timepoints, B), or 2-way ANOVA, with overall comparisons indicated on the graphs (C-F), ****P<0.0001.

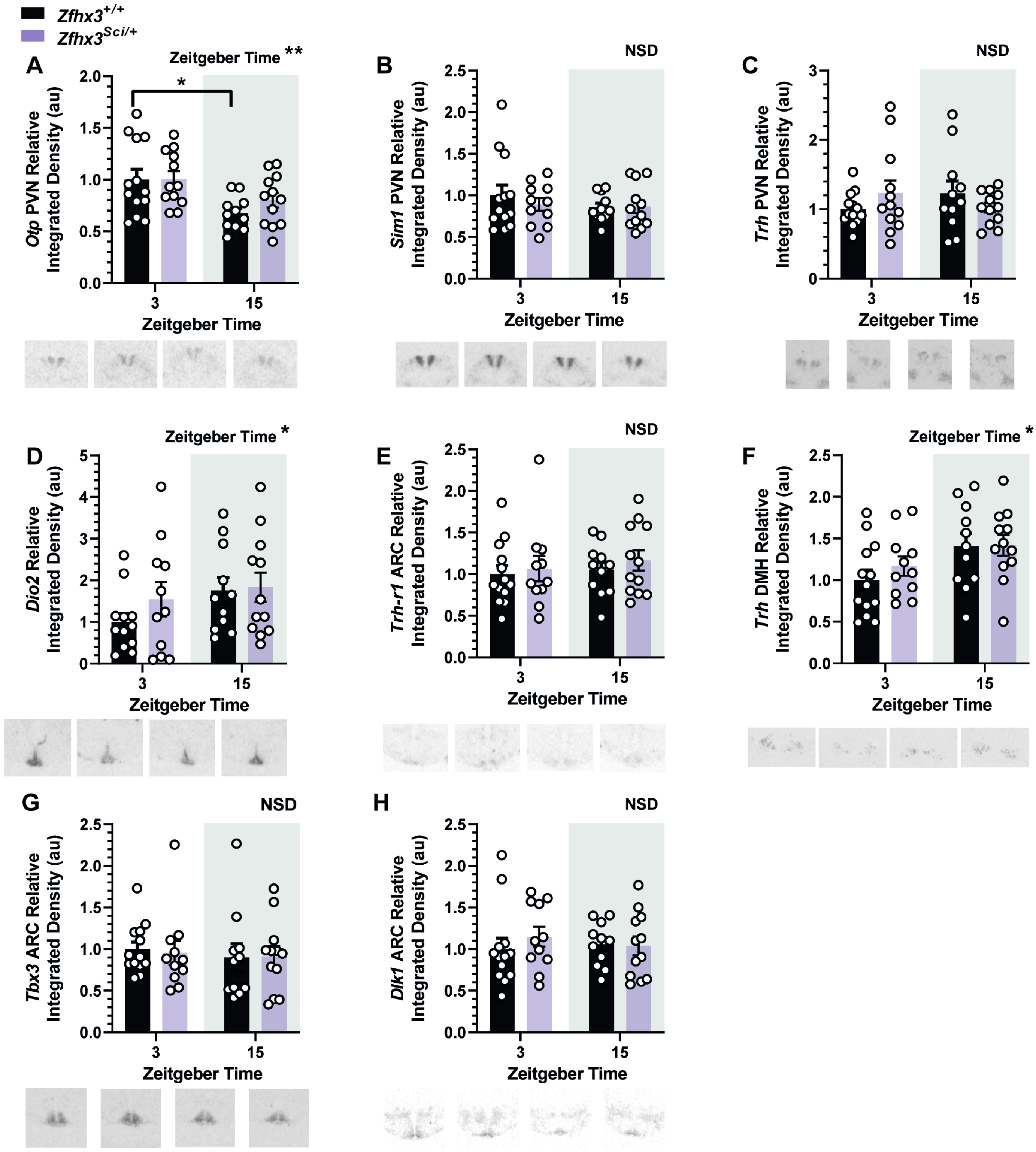


**Supplemental figure 3: Hypothalamic candidate gene expression not altered in female *Zfhx3*$^{Sci/+}$ mice**

Expression of *Otp* in the paraventricular nucleus (PVN) was reduced at ZT 15 (A), *Sim1* (B) and *Trh* (B) expression in the PVN were not altered by genotype or time. Expression of *Dio2* in the ventricular ependymal layer increased overall at ZT 15 (D) *Trh-r1* expression in the ARC was unaltered by time or genotype (E), while *Trh* expression in the dorsomedial hypothalamus was increased at ZT 15 (F), *Tbx3* (G) and *Dlk1* (H) expression in the ARC were unaltered by time or genotype. Plotted are mean ± SEM with individual values overlaid. Example images are shown beneath each plot. Comparisons are by 2-way ANOVA with Šídák's multiple comparison tests indicated where appropriate. *Dio2* data were log transformed for statistical comparison. * $P < 0.05$, ** $P < 0.01$, NSD: no significant differences. N = 10 – 12.
